# Cancer cells surviving cisplatin chemotherapy increase stress-induced OMA1 activity and mitochondrial fragmentation

**DOI:** 10.1101/2025.09.29.679325

**Authors:** Melvin Li, Chenille A. McCullum, Louis T.A. Rolle, Qin Ni, Zhuoxu Ge, Sean X. Sun, Kenneth J. Pienta, Sarah R. Amend

## Abstract

Cancer is one of the leading causes of deaths worldwide. Once cancer cells acquire therapy resistance, they become the main driver of cancer lethality in patients. Thus, mechanisms of therapy resistance must be investigated to improve patient outcomes. Mitochondria are critical organelles in the cellular stress responses, undergoing dynamic morphological and functional changes in response to external stimuli. We and others have identified a chemotherapy-resistant cancer cell state where cells that survive treatment exhibit a dramatic increase in cell size and remain non-proliferative for weeks. In this study, we demonstrate that cancer cells that enter this resistant cell state in response to cisplatin increase OMA1 activity and decrease mitochondrial fusion and function to combat oxidative stress. These findings contribute to further understanding the role of the mitochondrial stress responses in therapy resistance in cancer and provide a potential therapeutic avenue to targeting cancer cells that enter this chemotherapy-resistant cell state.

## INTRODUCTION

Mitochondria are essential organelles that play multi-faceted roles in maintaining cellular homeostasis and stress responses^1,2^. These organelles undergo dynamic morphological changes in response to metabolic and stress stimuli^1,2^. Mitochondrial fission is induced by oxidative stress, resulting in a fragmented phenotype to decrease oxidative capacity or to shuttle damaged mitochondria for degradation via mitophagy^3^. Mitochondrial fusion can be induced by increased energetic demands, creating a hyperfused branching network with increased oxidative capacity^3^. These organelles are dysregulated in many cancers, enabling cells to better obtain resources and adapt to their environment^3^. Recent studies have shown that alterations in mitochondrial morphology and dynamics confer resistance to a wide range of chemotherapies across many tumor types^4–9^. Given the evidence on the role of mitochondria in therapy-resistant cancers, targeting mitochondria is a promising strategy to combat cancer drug resistance.

Our group and others have previously reported that cancer cells that survive chemotherapy enter a resistant cell state in which they increase in size over time and are resistant to subsequent treatments^10–14^. This phenomenon has been reported in mouse models and in human tumors across various tumor types and is induced by multiple classes of chemotherapy and environmental conditions^13–19^. In addition, the presence of this cell state in patient primary tumors predict progression to metastatic disease, further strengthening its significance *in vivo*^13,20,21^. Our previous work has identified that these cells exhibit altered nuclear morphology, a dramatic increase and displacement of intracellular labile iron and lysosomes, and increased antioxidant responses through NRF2 signaling^10,22^. Mitochondria interact with all these processes, including communicating with the nucleus through retrograde signaling, storing labile iron in iron-sulfur clusters, colocalizing with lysosomes to degrade damaged portions of mitochondria, and undergoing turnover in response to NRF2 signaling^23–28^.

Mitochondrial mass has been reported to increase in cancer cells entering this resistant state from paclitaxel treatment in breast cancer, while another study observed increased mitophagy in cancer cells surviving chemotherapy treatment in a head and neck cancer model^29,30^. Beyond these observations, the molecular mechanisms of the mitochondrial phenotypes in this chemotherapy-resistant cancer cell state have been largely unexplored. In this study, we investigated mitochondrial morphology, dynamics, and stress responses in prostate cancer cells surviving cisplatin chemotherapy. Through various imaging modalities and protein expression analyses, we identified that the cells surviving chemotherapy exhibit increased OMA1 activity, aberrant cristae morphology, and reduced mitochondrial fusion, leading to decreased mitochondrial function. Our findings suggest that targeting OMA1 activity could be a potential therapeutic vulnerability specific to this resistant phenotype.

## RESULTS

### Cancer cells surviving chemotherapy exhibit increased levels of reactive oxygen species and increased mitochondrial fragmentation

The PC3 prostate cancer cell line was treated with an LD50 dose of cisplatin (6 µM) for 72 hours^10,31^. At the end of treatment, cisplatin-containing media was removed, and the surviving cells were then cultured in fresh media for up to 10 days (**Fig. 1a**). During the 10-day period after cisplatin removal, surviving cells were not proliferative and increased in cell size over time, resulting in a 40-fold increase in cell volume at 10 Days Post-Treatment Removal (10 Days PTR) (**Fig. 1b-c**). We have previously published that cells continue to die via apoptosis after treatment removal, and the death rate plateaus at 10 Days PTR^10^. We initially hypothesized that these surviving cells have recovered from the cellular damage induced by chemotherapy, including the increased oxidative stress and DNA damage induced by chemotherapy treatment^32^. We found that the cells 10 Days PTR exhibited increased levels of reactive oxygen species (ROS) per cell area when compared to untreated cells (**Fig. 1d-e**). To validate this finding, both groups were treated with antioxidant N-acetyl cystine (NAC) for 24 hours. While the PC3 cells did not show any change in DCF-DA staining, cells 10 Days PTR exhibited a decrease in DCF-DA staining under NAC-treated conditions, (**Supplementary Fig. S1**).

**Fig. 1.**
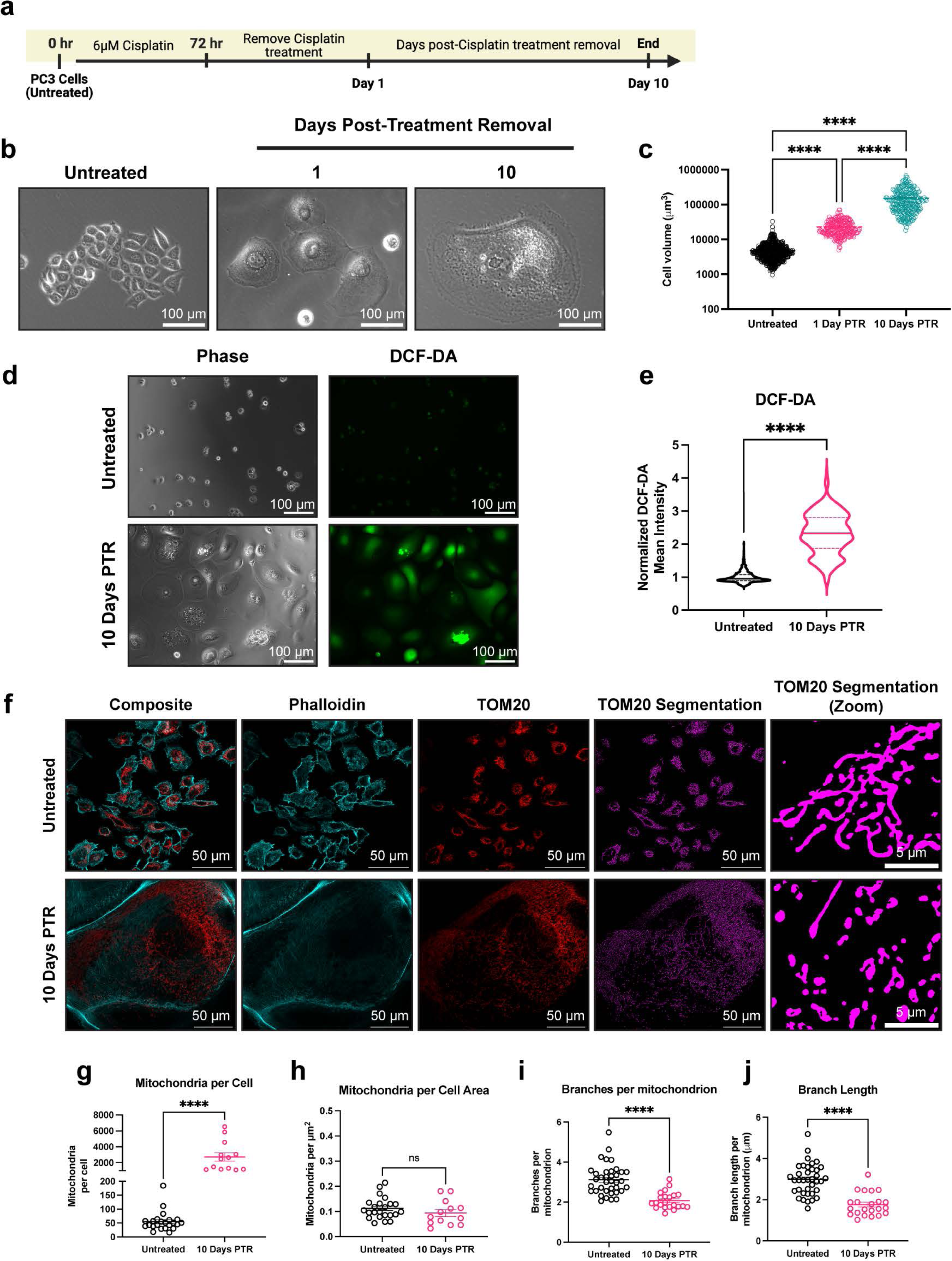
Cancer cells surviving chemotherapy have increased levels of reactive oxygen species and exhibit mitochondrial fragmentation. (**a**) Timeline for the induction of the chemotherapy-resistant cancer cell state. Created in BioRender. Li, M. (2025) https://BioRender.com/vkhgx05 (**b**) Phase contrast images of untreated cells and cells 1- and 10-Days Post-Treatment Removal (PTR). (**c**) Cell volume measurements of untreated cells and cells 1- and 10-Days PTR. (**d**) Phase contrast images and DCF-DA fluorescence images in untreated cells and cells 10 Days PTR. (**e**) Quantification of mean fluorescence intensity of DCF-DA staining. (**f**) Representative images of TOM20 and Phalloidin in untreated cells and cells 10 Days PTR. Segmentation of TOM20 signal was performed to quantify mitochondrial morphology features. (**g**) Quantification of number of mitochondria per cell. (**h**) Quantification of mitochondria normalized to cell area. (**i**) Quantification of number of branches per mitochondrion. (**j**) Quantification of branch length per mitochondrion. n = 679, 257, and 249 cells for untreated, cells 1 Day PTR, and cells 10 Days PTR, respectively (**c**); n = 1141 and 362 cells for untreated and cells 10 Days PTR, respectively (**e**); n = 37 and 22 cells for untreated and cells 10 Days PTR, respectively (**g-j**). Data are presented as mean ± s.e.m. (**c**, **g-j**), while data in violin plots were presented as median and corresponding interquartile ranges (**e**). p values were calculated with a one-way ANOVA followed by a post-hoc Tukey’s multiple comparisons test (**c**), a two-tailed Mann Whitney test (**e**), and an unpaired Student’s two-tailed t-test (**g-j**). ns not significant, **** p<0.0001.

Mitochondrial morphology is altered under oxidative stress^33^. We observed that cells 10 Days PTR have more mitochondria on a per cell basis, although this difference is non-significant when normalizing to cell area (**Fig. 1f-h**). Quantification of mitochondrial morphology showed that cells 10 Days PTR exhibited mitochondrial fragmentation, with a 1.5-fold decrease in branches per mitochondrion and a 1.5-fold decrease in branch length per mitochondrion (**Fig. 1f, i-j**).

### Surviving cells increase DRP1 localization to mitochondria

Mitochondrial fragmentation can be the result of an increase in fission or a decrease in fusion^34^. We probed for the expression of dynamin-related GTPases that are known to remodel mitochondrial structure. Dynamin-related protein 1 (DRP1) is the main driver of mitochondrial fission at the outer mitochondrial membrane (OMM) upon translocation from the cytosol to the mitochondria. Phosphorylation of DRP1 at Ser616 promotes its translocation to the OMM, while phosphorylation of DRP1 at Ser637 prevents the translocation from occurring^1^. Total DRP1 expression was unchanged in cells 10 days PTR compared to control and Ser616 phosphorylation was not detected in either group (**Fig. 2a**). Cells 10 Days PTR had decreased DRP1 Ser637 phosphorylation compared to untreated cells, suggesting a decrease in DRP1 inhibition and an increase in DRP1 localization to mitochondria (**Fig. 2a**). DRP1 binds to receptors mitochondrial fission factor (MFF), MiD49, or MiD51 on the OMM that allow DRP1 to assemble on the mitochondrial surface for its fission activity^1,2,35^. Compared to control, cells 10 Days PTR showed increased expression of MFF and MiD49, but a decrease in MiD51 expression (**Fig. 2a**). We next assessed DRP1 mitochondrial localization via immunofluorescence. Cells 10 Days PTR had increased DRP1-mitochondria localization when compared to PC3 control cells (**Fig. 2b-f**). These results suggest that DRP1 may play a role in the fragmentation phenotype that we observe in the mitochondria of surviving cells.

**Fig. 2.**
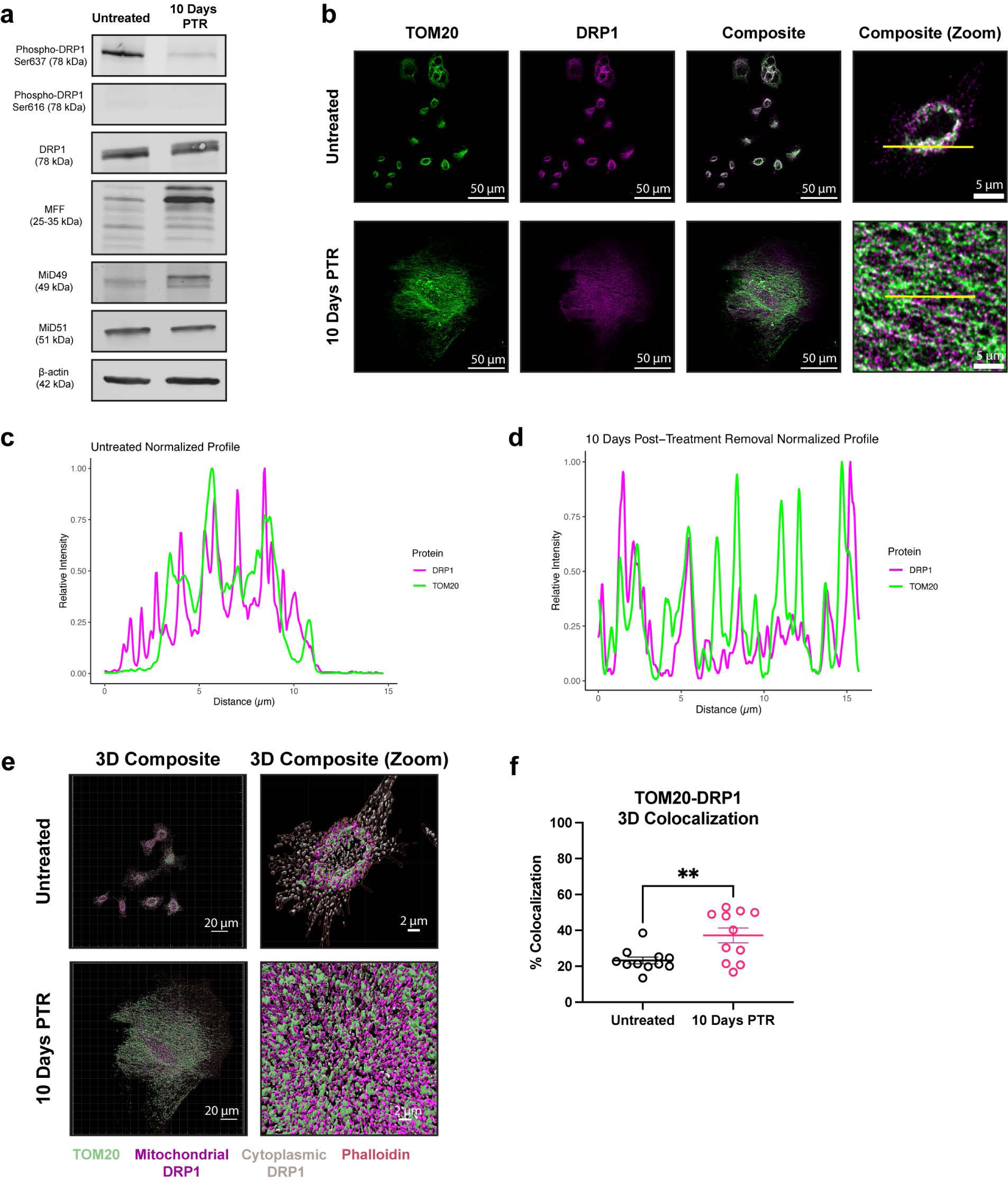
Cells 10 Days PTR increase DRP1 localization to mitochondria. (**a**) Representative western blots of Phospho-DRP1 Ser637, Phospho-DRP1 Ser616, DRP1, MFF, MiD49, MiD51, and β-actin loading control. n = 3 biological replicates. We note that due to similarities in molecular weight and identical primary antibody species, each protein was probed independently on separate blots with their own separate β-actin loading control. Original, uncropped blots (including the β-actin loading control for each blot) are presented in Supplementary Figure S2. (**b**) Max intensity projection images of TOM20 and DRP1 channels in untreated cells and cells 10 Days PTR. (**c**) Normalized profile intensity plot of TOM20 and DRP1 signal in untreated cells. (**d**) Normalized profile intensity plot of TOM20 and DRP1 signal in cells 10 Days PTR. (**e**) 3-dimensional (3D) rendering of TOM20, DRP1, and Phalloidin signal in untreated cells and cells 10 Days PTR. (**f**) Quantification of TOM20-DRP1 colocalization in 3D. n = 11 cells each for untreated and cells 10 Days PTR (**f**). Data are presented as mean ± s.e.m. in (**f**), and p values were calculated with an unpaired Student’s two-tailed t-test (**f**). ** p<0.01.

### Mitochondria in surviving cells exhibit aberrant cristae morphology

In addition to DRP1-mediated fission, mitochondrial fragmentation can occur through a decrease in fusion, which is mainly regulated by Mitofusin 1 (MFN1), Mitofusin 2 (MFN2), and optic atrophy 1 protein (OPA1) ^1,35^. MFN1 and MFN2 are responsible for the fusion of the OMM ^1^. There were no differences in MFN1 and MFN2 expression between surviving cells and untreated cells (**Fig. 3a**). OPA1 is an inner membrane remodeler that maintains cristae structure^1,35^. The long isoforms of OPA1 mediate fusion of the inner mitochondrial membrane and promote tight stacking of lamellar cristae^36,37^. Under oxidative stress or mitochondrial depolarization, OPA1 is cleaved by OMA1 into short isoforms (L1 to S3, and L2 to S5), leading to decreases in mitochondrial fusion and energetic capacity^37–39^.

**Fig. 3.**
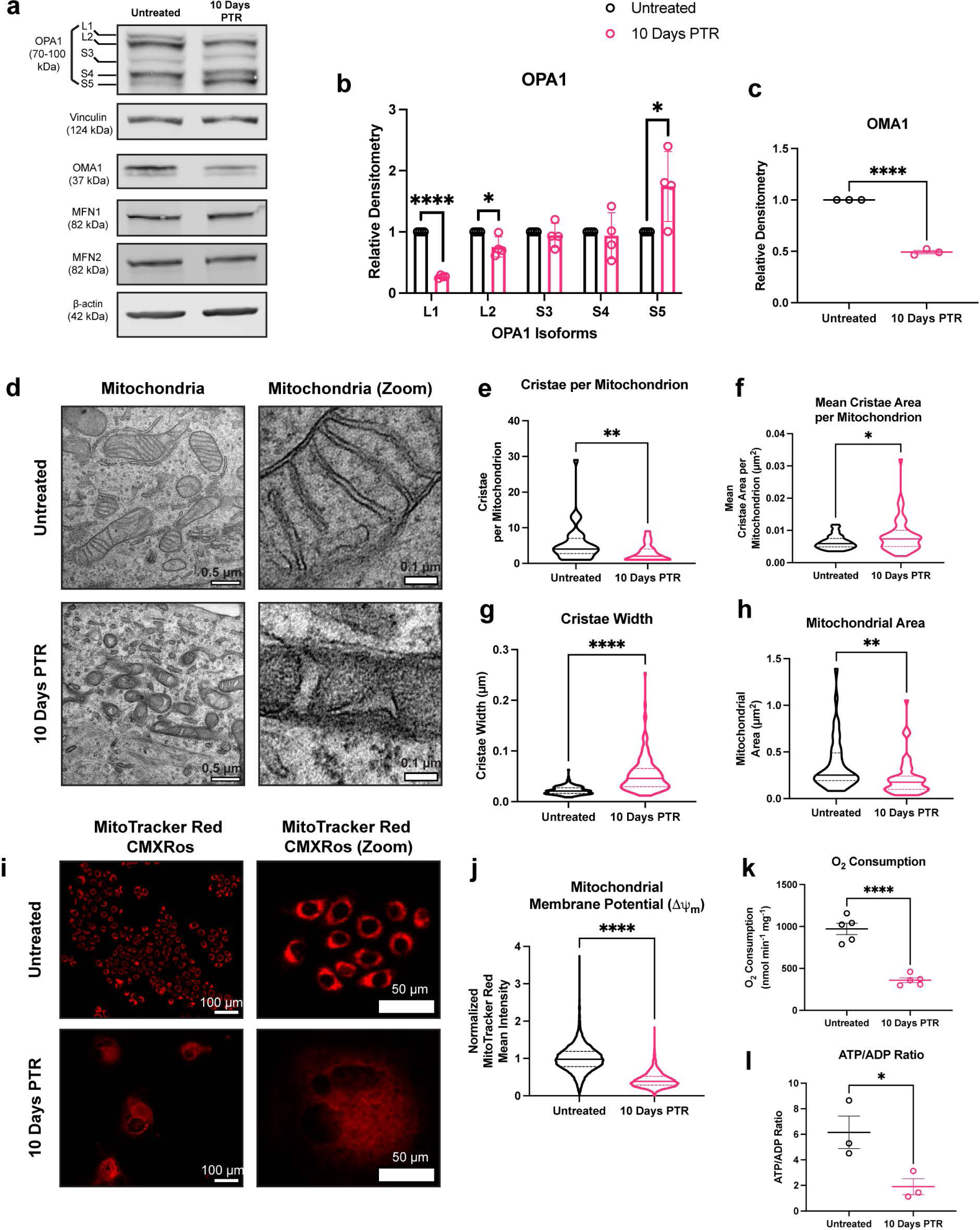
Mitochondria in cells 10 Days PTR have aberrant cristae morphology. (**a**) Representative western blots of OPA1, OMA1, Vinculin, and β-actin loading control. We note that due to similarities in molecular weight and identical primary antibody species, each protein was probed independently on separate blots with their own separate β-actin or vinculin loading control. Original, uncropped blots (including the β-actin or vinculin loading control for each blot) are presented in Supplementary Figure S3. **b**) Quantifications of OPA1 isoform protein expression in untreated cells and cells 10 Days PTR. Samples derive from the same experiment in each blot that was quantified. (**c**) Quantification of OMA1 expression in untreated cells and cells 10 Days PTR. Samples derive from the same experiment in each blot that was quantified. (**d**) Representative transmission electron microscopy (TEM) images of mitochondria in untreated cells and cells 10 Days PTR. (**e**) Quantification of number of cristae per mitochondrion. (**f**) Quantification of mean cristae area per mitochondrion. (**g**) Quantification of cristae width. (**h**) Quantification of mitochondrial area between untreated cells and cells 10 Days PTR. (**i**) Representative fluorescence images of MitoTracker Red CMXRos staining in untreated cells and cells 10 Days PTR. (**j**) Quantification of mean fluorescence intensity of MitoTracker Red CMXRos staining between indicated groups. (**k**) Oxygen consumption rate normalized to milligram of cell weight in untreated cells and cells 10 Days PTR. (**l**) Quantification of ATP/ADP ratio in untreated cells and cells 10 Days PTR. n = 4 biological replicates (**b**); n = 3 biological replicates (**c**); n = 42 and 59 mitochondria for untreated and cells 10 Days PTR, respectively (**e-f, h**); n = 229 and 177 cristae for untreated and cells 10 Days PTR, respectively (**g**); n = 12,147 and 2502 cells for untreated and cells 10 Days PTR, respectively (**j**). n = 5 biological replicates (**k**); n = 3 biological replicates (**l**). Data are presented as mean ± s.e.m. (**b-c**), while data in violin plots are presented as median and corresponding interquartile ranges (**e-h, j**). p values were calculated with a two-tailed Kolmogorov-Smirnov test (**e-h**), a two-tailed Mann-Whitney test (**j**), and two-tailed unpaired t-test (**k-l**). * p<0.05, ** p<0.01, **** p<0.0001.

Cells that survived cisplatin treatment showed a decrease in the L2 isoform of OPA1 and an increase of the S5 isoform, suggesting that L-OPA1 is cleaved to S-OPA1 by OMA1 (**Fig. 3a-b**). Upon stress conditions, OMA1 cleaves itself when activated to act on L-OPA1^39,40^. We observed decreased expression of OMA1 in cells 10 Days PTR when compared to PC3 control (**Fig. 3a, c**). Taken together, the altered levels of L2 and S5 isoforms of OPA1 and decreased OMA1 levels in cells 10 Days PTR suggest OMA1 activity is higher in the cells that survive cisplatin than in control cells.

OPA1 is responsible for maintaining mitochondrial cristae structure. To evaluate this directly, we performed transmission electron microscopy (TEM) to assess mitochondrial cristae morphology. Mitochondria in untreated cells had tight lamellar cristae, while the mitochondria in cells 10 Days PTR exhibited amorphous and wider cristae shape (**Fig. 3d**). Quantification of cristae morphology showed that mitochondria in cells 10 Days PTR had fewer cristae, higher mean cristae area, and higher cristae width (**Fig. 3e-g**). In addition, mitochondria in cells 10 Days PTR were 3.5 times smaller by area than the mitochondria in untreated cells, supporting the fragmentation phenotype that we observed in the immunofluorescence imaging in **Fig. 1f, i-j** (**Fig. 3h**).

Overall, these data indicate that mitochondria in cells 10 Days PTR have impaired cristae organization. The complexes of the electron transport chain (ETC) reside on the cristae structure. The ETC complexes generate a negative membrane potential (ΔΨ_m_) across the inner mitochondrial membrane (IMM) as electrons are shuttled through^41^. To assess the functional implications of this altered mitochondrial cristae structure, we measured the mitochondrial membrane potential via live-cell staining with MitoTracker Red CMXRos. We observed that the mitochondria in cells 10 Days PTR had a 2.5-fold decrease in MitoTracker Red CMXRos signal when compared to untreated cells, indicating that mitochondria are less active in cells 10 Days PTR (**Fig. 3i-j**). We then took oxygen consumption measurements over time as an orthogonal method to validate our findings. The oxygen consumption rate (OCR) of cells 10 Days PTR was 2.7-fold lower than untreated cells (**Fig. 3k**). Given that mitochondria are a major source of ATP generation, we evaluated the energetic status of the cells through the ratio of ATP relative to ADP and observed that cells 10 Days PTR had a 3-fold decrease in ATP/ADP ratio when compared to the untreated cells (**Fig. 3l**) ^42^. The decrease in ATP/ADP in cells 10 Days PTR could be a result of a shift from mitochondrial metabolism to glycolysis. This hypothesis is supported through the enrichment of HALLMARK GLYCOLYSIS and HALLMARK HYPOXIA gene sets via gene set enrichment analysis (GSEA) from our single-cell RNA sequencing data (**Supplementary Fig. S2a-b**). Taken together, these data suggest that mitochondria in cells 10 Days PTR display aberrant cristae morphology and have decreased functional output when compared to mitochondria in untreated cells.

### Cells that survive cisplatin treatment decrease both mitochondrial fission and fusion dynamics

Given the changes in both DRP1 phosphorylation and OPA1 processing in cells 10 Days PTR, we next directly evaluated mitochondrial fission and fusion dynamics. We performed time lapse imaging to capture mitochondrial fission and fusion events on a per-mitochondrion basis, taking an image every 30 seconds for a total of 15 minutes. Overall, we observed that the fission and fusion rate scaled with increasing mitochondrial area (**Fig. 4a-c**). When normalized to mitochondrial area, cells 10 Days PTR had a 50% decrease in fission rate and a 50% decrease in fusion rate than untreated cells (**Fig. 4a, d-e, Supplementary Video 1-4**).

**Fig. 4.**
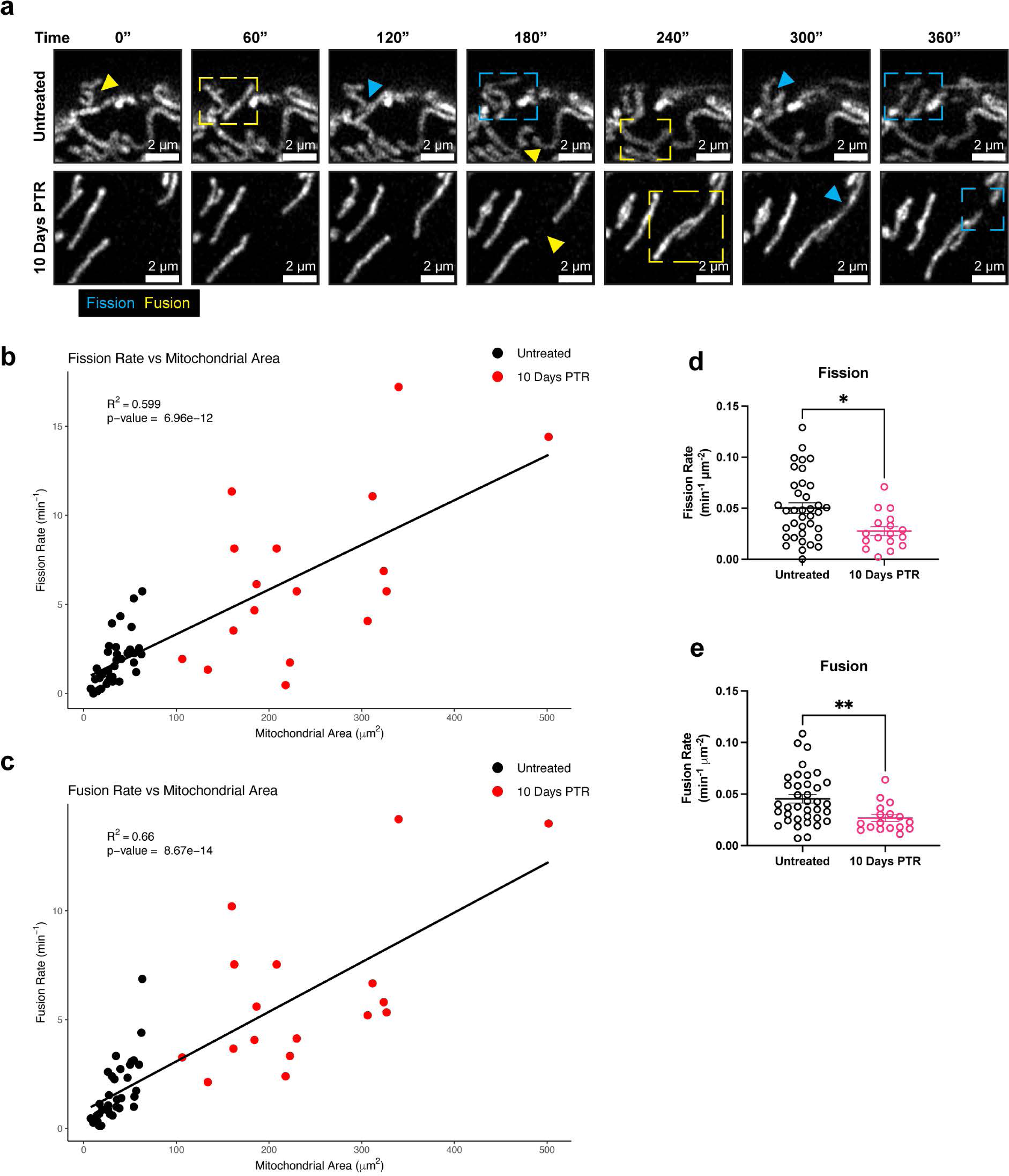
Cells 10 Days PTR decrease both mitochondrial fission and fusion dynamics. (**a**) Representative fluorescence images of mitochondria every 60 seconds for 6 minutes in untreated cells and cells 10 Days PTR. Yellow arrow and box indicate a fusion event, while a blue arrow and box indicate a fission event. (**b**) Linear regression analysis of fission rate vs mitochondrial area. (**c**) Linear regression analysis of fusion rate vs mitochondrial area. (**d**) Quantification of fission rate in untreated cells and cells 10 Days PTR when normalized to mitochondrial area. (**e**) Quantification of fusion rate in untreated cells and cells 10 Days PTR when normalized to mitochondrial area. n = 37 and 17 cells for untreated and cells 10 Days PTR, respectively (**b-c, d-e**). Data are presented as scatter plots (**b-c**) and mean ± s.e.m. (**d-e**). R-squared values were calculated with the lm() and summary() functions in RStudio^93^ (**b-c**). p values were calculated with the lm() and summary() functions in RStudio^93^ (**b-c**) and a two-tailed Mann-Whitney test (**d-e**). * p<0.05, ** p<0.01.

### *Oma1* knockdown promotes cell death in cancer cells surviving chemotherapy

Given that we observed decreases in mitochondrial fusion rather than an increase in mitochondrial fission in cells 10 Days PTR from the time lapse experiments, we hypothesized that the changes in OPA1 processing through OMA1 play a critical role in the mitochondrial response against oxidative stress in these cells. To test this, we knocked down *Oma1* using siRNA and observed changes in OMA1 protein expression. In untreated cells, OMA1 was knocked down with 80% efficiency, while there was only a 50% knockdown of OMA1 in cells 10 Days PTR (**Fig. 5a-c**). In untreated cells, one of the *Oma1* siRNAs increased cell viability over the scramble negative control while the other had no effect (**Fig. 5d-e**). *Oma1* knockdown resulted in a 20-40% decrease in cell viability in cells 10 Days PTR when compared to the scramble negative control, suggesting a modest susceptibility of cells 10 Days PTR to *Oma1* silencing (**Fig. 5d, f**).

**Fig. 5.**
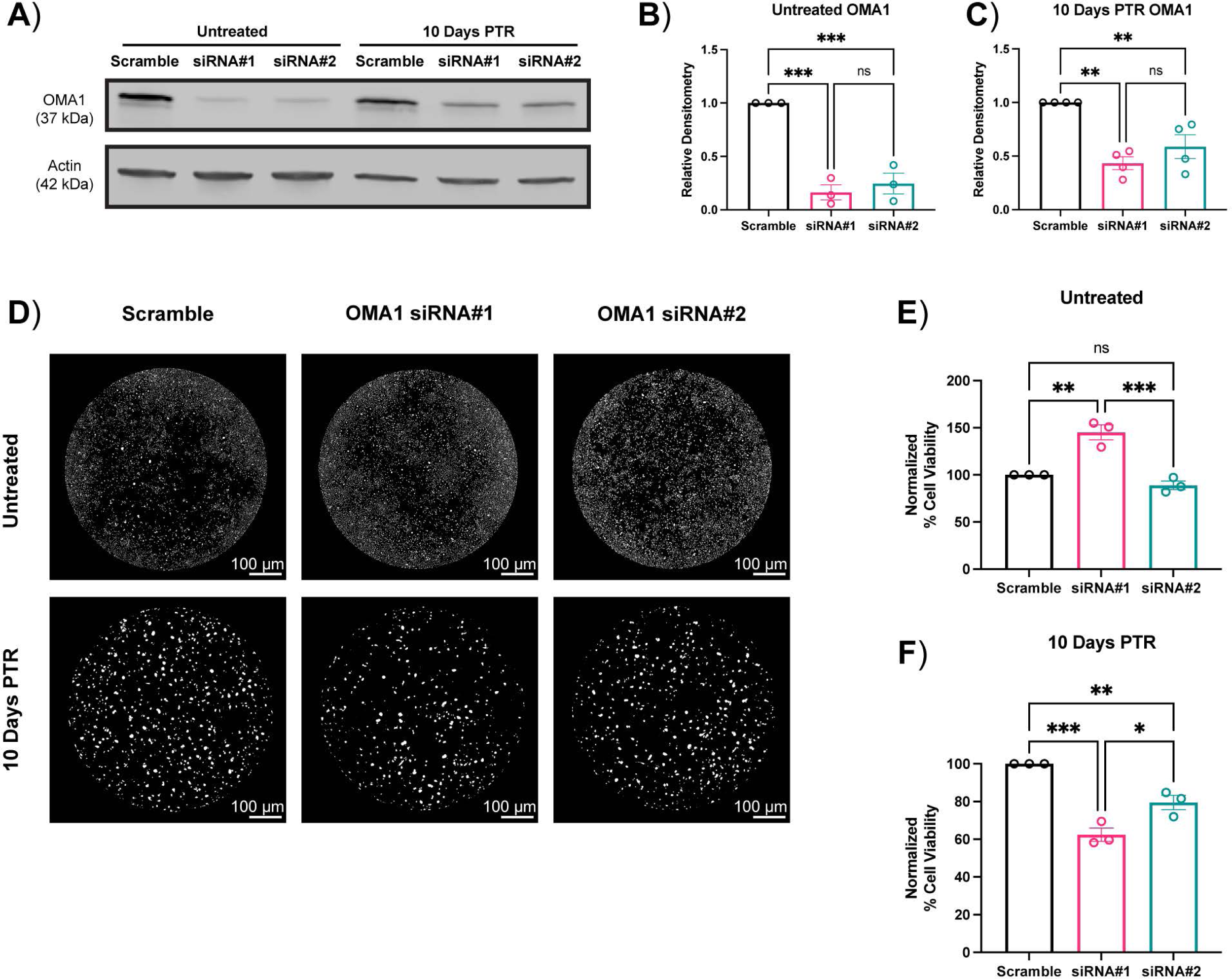
*Oma1* knockdown promotes cell death in cancer cells surviving chemotherapy. (**a**) Representative western blot of OMA1 after 72 hours of siRNA or Scramble transfection in untreated cells and cells 10 Days PTR. (**b**) Quantification of OMA1 protein expression in untreated cells after 72 hours of siRNA or Scramble transfection. (**c**) Quantification of OMA1 protein expression in cells 10 Days PTR after 72 hours of siRNA or Scramble transfection. (**d**) Representative images of one well of a 96-well plate of Hoechst-stained nuclei from untreated cells and cells 10 Days PTR after 72 hours of siRNA or Scramble transfection. (**e**) Quantification of cell viability normalized to scramble negative control in untreated cells. (**f**) Quantification of cell viability normalized to scramble negative control in cells 10 Days PTR. n = 3 biological replicates (**b, e-f**); n = 4 biological replicates (**c**). Data are presented as mean ± s.e.m. (**b-c, e-f**). p values were calculated with a two-tailed unpaired t-test (**b-c, e-f**). ns not significant, * p<0.05, ** p<0.01, *** p<0.001.

## DISCUSSION

Given the critical roles of mitochondria in cellular stress responses and their implications in therapy-resistant cancer cells, recent advancements in identifying molecular regulators that drive morphological and functional adaptations provide potential therapeutic avenues for cancer^3,4,6,35,43^. Our group and others have reported the emergence of an enlarged and non-proliferative cancer cell state in response to various chemotherapy treatments ^10–14,22^. Cells continued to undergo apoptosis after treatment was removed, with the death rate decreasing over time and plateauing by 10 Days PTR^10^. These cells are resistant to subsequent chemotherapy and are associated with worse prognosis in mouse models and in patients^13–20^. Morphologically, cells entering this resistant state resemble those the phenotype of therapy-induced senescent cells. While the cancer cells surviving chemotherapy in our model exhibit some canonical features of senescent cells, such as an increase in cell size and sustained DNA damage, work from our group has shown that they continue to progress through the cell cycle without cell division^10,44–47^. Whether cells in this resistant state and therapy-induced senescent cells represent the same phenotype remains an active area of research^48–52^. In this study, we evaluated the mitochondrial stress responses in cancer cells that survive in the days following chemotherapy.

Cells 10 Days PTR had increased levels of ROS (**Fig. 1d-e**), consistent with previous reports in bladder cancer where giant cells that emerge after cisplatin treatment survive with increased levels of ROS^53^. The accumulation of ROS can induce oxidation of macromolecules that are essential for cellular homeostasis, leading to DNA damage, misfolding of proteins, and lipid peroxidation^54–57^. We have previously published that cells surviving chemotherapy have continued DNA damage response (DDR), even at 10- and 15-Days PTR^10^, and that cells 10 Days PTR exhibit increased lipid peroxidation^22^. Moreover, this work also showed that these surviving cells have an increase in nuclear factor erythroid 2-related factor (NRF2)-mediated antioxidant response, further demonstrating that they are in a state of oxidative stress^22^.

Mitochondrial biogenesis, morphology, and function are altered under oxidative stress^33,58–61^. While we observe that surviving cells have more mitochondria on a per cell basis, when normalizing to cell size, there was no difference in mitochondrial number (**Fig. 1f-h**). When analyzing the structure of the mitochondria, we found that the mitochondria in cells 10 Days PTR were more fragmented and had fewer branches per mitochondrion when compared to the mitochondria of the untreated cells (**Fig. 1f, i-j**). We hypothesize that mitochondria in surviving cells underwent fragmentation in response to the increase in ROS levels when compared to untreated cells. Building on observations of this fragmentation phenotype in various cell types and cancer cell lines subjected to oxidative stress^5,8,58,60,62–64^, our findings align with recent work by Song et al. showing that esophageal squamous cell carcinoma cancer cells with a similar morphological phenotype that survive paclitaxel treatment had more mitochondria per cell but exhibited a smaller and more fragmented phenotype when compared to the untreated controls^65^. The mitochondria in those surviving cells also moved more slowly and had fewer fission and fusion events than the mitochondria in the control cells^65^. This suggests that these treatment-induced mitochondrial phenotypes could span across different cancer types and across therapies that act through different mechanisms, as paclitaxel is a microtubule stabilizer, while cisplatin creates adducts in purine bases and ultimately causes DNA double strand breaks^66,67^.

When probing for the expression of the molecular regulators of mitochondrial fusion, we identified differential expression of multiple isoforms of OPA1 when comparing cells 10 Days PTR and untreated cells (**Fig. 3a-b**). The decrease in L2-OPA1 and increase in S5-OPA1 expression suggested that the stress-induced protease OMA1 was active in cells 10 Days PTR (**Fig. 3a-b**). This is further supported by the decreased level of OMA1 itself, likely due to its autocatalytic turnover. The S3 and S5 isoforms of OPA1 are fusion incompetent, and have been shown to induce mitochondrial fission through interactions with DRP1 and other fission machinery and were shown to be protective in events of oxidative stress^38,68^. The accumulation of these OPA1 isoforms may be a mechanism to reduce mitochondrial fusion and decrease OXPHOS to prevent further production of ROS, as mitochondria are major sources of intracellular ROS when electrons are shuttled through the ETC^59^.

We observed that the mitochondria in cells 10 Days PTR had larger and more amorphous cristae than the tight, lamellar cristae in the mitochondria of untreated cells (**Fig. 3d-h**). This aberrant morphology can be attributed to the decrease in L-OPA1 and accumulation of S-OPA1 via OMA1 cleavage (**Fig. 3a-c**), as the short isoforms are not tethered to the IMM to promote cristae folding^36,39,40^. Cristae folding increases the surface area of the IMM for oxidative phosphorylation by bringing electron transport chain (ETC) complexes closer together and concentrating the proton gradient^36,69,70^. Cells 10 Days PTR had decreased ETC efficiency based on MitoTracker Red CMXRos staining, along with reduced oxygen consumption rate and ATP/ADP ratios, collectively indicating decreased mitochondrial function (**Fig. 3i-l**). It has been reported that cells engineered to express S-OPA1 exclusively have higher survival under oxidative stress than cells engineered to express only L-OPA1^68^. Cells expressing solely L-OPA1 produced more superoxide radicals and therefore were sensitized to oxidative stress^68^.

Thus, the decrease in cristae folding in surviving cells further supports reduced OXPHOS as a survival mechanism under oxidative stress. Mitochondrial morphology is mediated by dynamic fission and fusion processes. We observed a surprising decrease in the mitochondrial fission rate in cells 10 Days PTR (**Fig. 4a, d**). This was unexpected given the parallel findings of increased DRP1 localization to mitochondria (via TOM20-DRP1 colocalization analysis), increased expression of MFF and MiD49, and decreased expression of phospho-DRP1 Ser637 (**Fig. 2a-f**). This may suggest an increase in the upstream signaling to promote mitochondrial fission, but also a mechanism downstream of DRP1 leading to the decreased fission rate we observed in the time lapse imaging. There are several factors that affect mitochondrial fission after DRP1 docks onto the various receptors on the OMM. The endoplasmic reticulum (ER) is a significant player in mitochondrial fission. The molecular regulators of mitochondrial fission and fusion colocalize with ER-mitochondrial contact sites to regulate mitochondrial dynamics^71,72^. The polymerization of actin around the ER and the mitochondria and the transfer of calcium from the ER to mitochondria mediate the constriction of the outer mitochondrial membrane that is required for mitochondrial fission to occur^73–76^. Cells 10 Days PTR may have decreased ER-mitochondrial contacts, which could explain the paradoxical increase in DRP1 localization to mitochondria and a decrease in mitochondrial fission rate, but this has yet to be tested. Based on the TEM images in **Fig. 3d**, the ER appear more fragmented and further from mitochondria in cells 10 Days PTR when compared to the ER in the untreated cells. Future work to validate these observations involves additional high-resolution microscopy studies to quantify ER morphology and ER-mitochondrial contact sites. In addition, investigating the role of the ER stress response pathway in surviving cells may provide more insight into the morphological remodeling we observe in the mitochondria^77,78^. Cytoskeletal filaments also play important roles in regulating mitochondrial fission, although their effects may be cell type or context specific. In a model of *X. laevis* melanocytes, depolymerizing microtubules with nocodazole decreased mitochondrial distribution, motility, and shape dynamics across the entire cell^79^, while mitochondrial associations with microtubules inhibited mitochondrial fission in a model of fission yeast^80^. How the cytoskeleton filaments affect mitochondrial dynamics in cancer cells in this resistant state is unknown and warrants further investigation. In addition to decreased mitochondrial fission rate, cells 10 Days PTR also exhibited a decrease in the rate of mitochondrial fusion (**Fig. 4a, e**). This data coincides with the increased OMA1 activity, decreased expression of the fusion-promoting L-OPA1 isoforms, and increased expression of the fusion-incompetent S-OPA1 isoforms we observed from **Fig. 3a-c**. The decrease in mitochondrial fusion could also lead to the fragmentation phenotype we observed in the mitochondria of the cells 10 Days PTR (**Fig. 1f, i-j**).

Based on our findings, we then hypothesized that OMA1 plays a critical role in the survival of cells 10 Days PTR. We transfected cells with *Oma1* siRNA and observed decreased OMA1 expression at the protein level in both groups (**Fig. 5a-c**). Cells 10 Days PTR exhibited a 20-40% decrease in cell viability after *Oma1* silencing while untreated cancer cells presented with an increase in cell viability or no change relative to scramble control (**Fig. 5d-f**). These results suggested that cells 10 Days PTR are modestly susceptible to *Oma1* silencing while untreated cancer cells are not. The variability in the cell viability response in untreated cells may be due to unknown off-target effects of siRNA#1, and further studies are needed to test this hypothesis. One limitation of this experiment was the use of transient siRNA to silence *Oma1*. While the *Oma1* siRNAs significantly reduced OMA1 protein expression in PC3 cells, they only achieved 50% knockdown efficiency in cells 10 Days PTR at the protein level (**Fig. 5a-c**). Thus, there may be some OMA1 activity retained in cells 10 Days PTR after *Oma1* siRNA transfection. The incomplete silencing of *Oma1* may have led to the modest decrease in cell viability in cells 10 Days PTR rather than a large decrease as we hypothesized (**Fig. 5a, c, d, f**). Future studies involve knocking out *Oma1* to ablate its expression at the protein level in PC3 cells and cells 10 Days and assessing its role in cancer cell survival after chemotherapy treatment removal. In addition, the functional implications of *Oma1* ablation in cells surviving chemotherapy such as changes in ROS levels and mitochondrial cristae organization should be investigated.

In summary, our findings suggest that cancer cells surviving chemotherapy increase OMA1 activity in response to oxidative stress. In this proposed pathway, stress-induced OMA1 activity cleaves OPA1 into its shorter isoforms, leading to a decrease in mitochondrial fusion, aberrant mitochondrial cristae morphology, and decreased efficiency of oxidative phosphorylation, ultimately promoting cell survival under increased ROS conditions (**Fig. 6**). Further validation of this mechanism is to be performed in future studies with rescue experiments such as probing OMA1 activity in surviving cells treated with the antioxidant N-acetyl cystine (NAC). Although there are currently no specific OMA1 inhibitors developed, this work contributes to the need for the screening of compounds against OMA1 activity, as other groups have solely used genetic mechanisms to inhibit OMA1 in various cancer models^43,81–83^. Given that OMA1 is predominantly dispensable in normal cells, likely due to functional redundancy of other mitochondrial proteases, inhibiting OMA1 is not expected to cause significant toxicity in healthy individuals. OMA1 deletion in various colorectal cancer mouse models reduced tumor progression and cristae re-organization without body weight loss or inflammation induction^83,84^. In addition, *Oma1*^-/-^ mice exhibited normal embryonic and adult development, presented with no changes in fertility, and did not experience any neurological pathologies when compared to wild type mice^85,86^. However, OMA1 deletion has been shown to exacerbate disease symptoms in certain pathological contexts, such as *Cox10*^-/-^-induced cardiomyopathy and diet-induced obesity^86,87^. To minimize on-target toxicities and maximize the therapeutic index, patients with pre-existing pathological conditions, such as obesity and cardiomyopathy, may need to be excluded from OMA1 targeting as an anti-cancer therapy in the clinical setting. Until specific OMA1 inhibitors are developed, further work involves knocking out *Oma1* in PC3 cells to assess its role in the entrance to and survival in this resistant cancer cell state. We hypothesize that these surviving cancer cells are critical mediator of cancer lethality in patients. Additional characterization of the cellular stress responses and molecular programs of cells in this resistant state will shine light on potential vulnerabilities that can be leveraged to target them for destruction and improve patient prognosis.

**Fig. 6.**
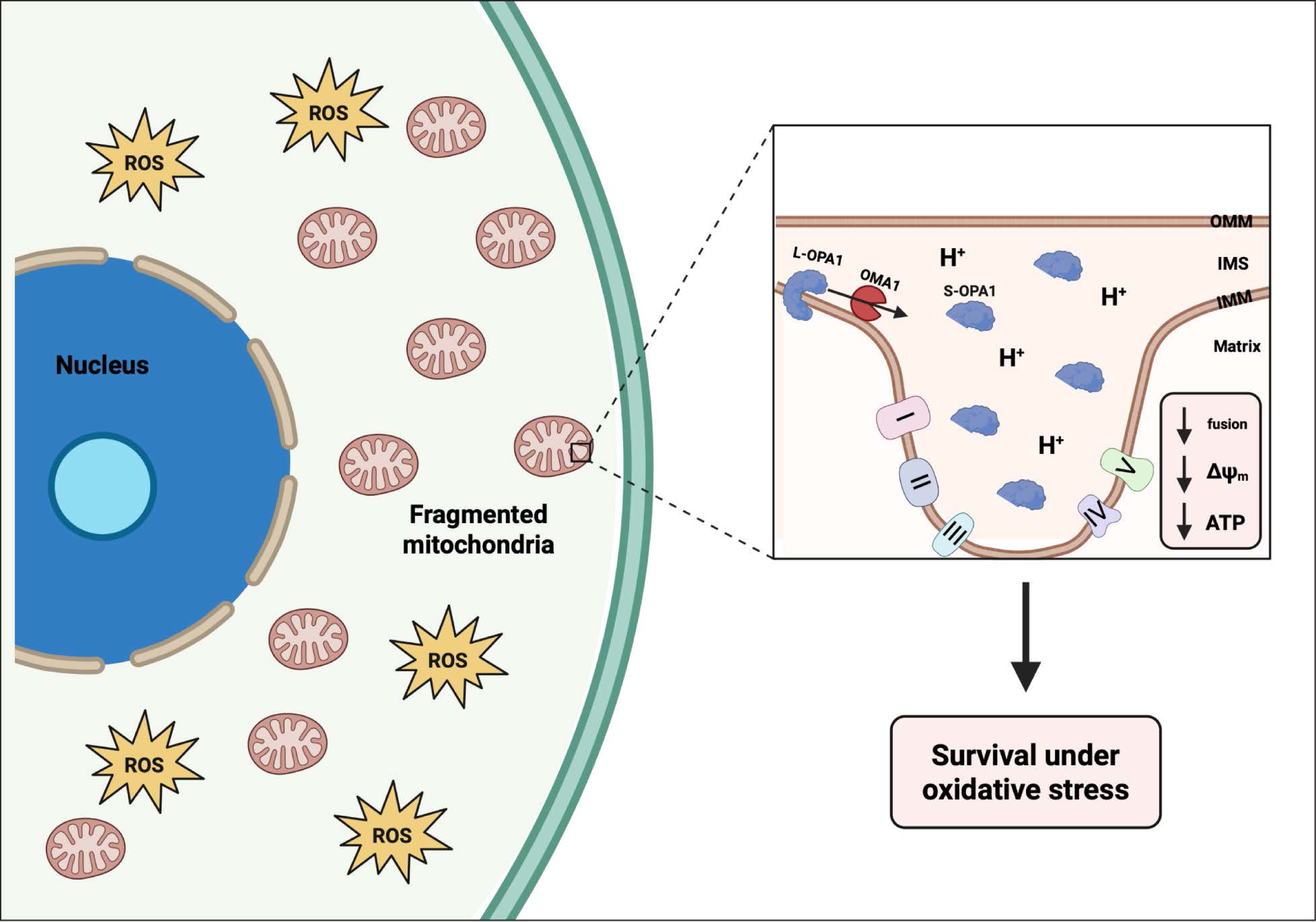
Cancer cells surviving chemotherapy increase OMA1 activity and decrease mitochondrial fusion and function to persist under oxidative stress. Cancer cells that survive chemotherapy enter a resistant cell state that increases their cell size and remain non-proliferative. Cells in this state have increased levels of reactive oxygen species (ROS) and exhibit fragmented mitochondria. Cells in this state increase OMA1 activity, cleaving L-OPA1 to S-OPA1, ultimately disrupting the mitochondrial cristae structure and reducing mitochondrial fusion. This altered cristae structure generates a diffuse proton gradient and decreases mitochondrial membrane potential, which lowers oxidative capacity as a survival mechanism against oxidative stress in these cells. This proposed mechanism requires further validation with rescue experiments in future studies. Created in BioRender. Li, M. (2025) https://BioRender.com/41jh49x.

## MATERIALS AND METHODS

### Cell culture

The PC3 cell line was purchased from American Type Culture Collection (ATCC). Cells were cultured in RPMI 1640 medium (ThermoScientific; Cat# 11875119) with 10% fetal bovine serum (FBS) (Avantar; Cat# 97068-085) and 1% penicillin and streptomycin (P/S) (Gibco; Cat#15140-122). Cells were incubated at 37℃ and 5% CO2.

### Induction of resistant cancer cell state

PC3 cells were plated in 150 mm dishes (Corning; Cat#: 353025) at a density of 1,250,000 cells per dish. 24 hours after seeding, cells were treated with an IC50 dose (6µM) of cisplatin (Millipore Sigma; Cat#: 232120) in RPMI media with 10% FBS and 1% P/S for 72 hours. Drug-containing media was then removed, and fresh cell culture media was added onto surviving cells. The surviving cells were monitored for 10 days, with media changes every 3 days. Cells were analyzed 10-days post-treatment removal (PTR) for all experiments.

### Phase contrast microscopy

Cells were cultured to their desired timepoint in T75 flasks. Phase contrast microscopy was performed on an EVOS M7000 (ThermoScientific) with a 10X objective.

### Cell volume measurement with Fluorescence eXclusion method

The microfluidic Fluorescence Exclusion method (FXm) was used to measure the volume of PC3 untreated cells and cells 10 Days PTR ^88,89^. In this method, microfluidic channels with known heights were fabricated. Cells were seeded into the device in the presence of a membrane-impermeable fluorescent dextran and imaged using an epifluorescence microscope. The reduction in fluorescence intensity due to cell volume exclusion is proportional to the ratio between the cell height and the channel height, allowing us to calculate cell volume. FXm channel fabrication and volume calculation are described in previously published work by Ni et al. and Rochman et al. ^89,90^.

### Immunofluorescence and Airyscan super-resolution imaging

Cells were plated onto poly-l-lysine coated coverslips (Thomas Scientific; Cat#: 1217N79). Cells were fixed with methanol-free 4% paraformaldehyde (ThermoFisher Scientific; Cat#28908) in 1X PBS for 15 minutes at room temperature followed by permeabilization with 0.2% Triton-X 100 (ThermoFisher Scientific; Cat#28314) in 1X PBS for 10 minutes at room temperature. Blocking was performed with 10% Normal Goat Serum (abcam; Cat# ab7481) in 0.1% PBS-Tween20 for 30 minutes at room temperature. Cells were incubated with primary antibody diluted in 10% Normal Goat Serum in 0.1% PBS-Tween20 overnight at 4℃: anti-Tom20 (Santa Cruz; Cat# sc-17764) diluted 1:100 and anti-DRP1 (Cell Signaling Technology; Cat#: 8570S) diluted 1:100. Secondary antibody incubation was performed using Goat anti-mouse IgG2a Cross-Absorbed Alexa Fluor 647 (ThermoFisher Scientific; Cat# A-21131) and Goat anti-rabbit IgG (H+L) Highly Cross-Absorbed Secondary Alexa Fluor Plus 555 diluted 1:1000 in 10% Normal Goat Serum in 0.1% PBS-Tween20 for 1 hour. Cells were also stained for F-actin during the secondary antibody incubation with Alexa Fluor 488 Phalloidin (ThermoFisher Scientific; Cat# A22287). Following secondary antibody and phalloidin incubation, cells were washed with 0.1% PBS-Tween20 three times and subsequently washed twice with 1X PBS. Coverslips were washed once in dH2O prior to mounting. Coverslips were mounted onto slides with 10 µL of Prolong Glass Antifade Mountant (ThermoFisher Scientific; Cat#: P36984) and samples were cured for at least 50 hours prior to imaging. All slides were imaged on a Zeiss LSM 880 microscope (Carl Zeiss) in Airyscan SR mode with an 63X, numerical aperture 1.4 PlanApo oil objective. Images were taken in a 3X3 tile z-stack at a resolution of 1768X1768 pixels per tile and an axial step size of 187 nm. 3D Airyscan Joint Deconvolution was performed on Zen software for image processing and downstream image analyses were performed in ImageJ^91^ and Imaris (Bitplane).

### Mitochondrial morphology analysis

Images were imported into ImageJ^91^ and converted to 8-bit images. The ImageJ pipeline, Mitochondria Analyzer, was used to analyze mitochondrial morphology^92^. In short, 3D thresholding was performed on each image within the Mitochondria Analyzer plugin^92^. The settings used were Rolling = 1.0, Max Radius = 1.25, Block Size = 3, C-Value = 3-5, Post-Processing Outlier Radius = 8.0 – 9.823 pixels. 2D slices were then obtained from the 3D thresholded images for 2D mitochondrial morphology analysis. Features such as branches per mitochondrion and branch length per mitochondrion were extracted in the Mitochondria Analyzer plugin with the 2D thresholded slices as input images.

### Colocalization analysis

In 2D, images were imported into ImageJ^91^ and split into individual channels and maximum Z-projections were generated for each channel. A line drawing was used to generate intensity profiles along the length of the line. Intensities were normalized within the DRP1 and Tom20 channel and overlayed together into a graph using RStudio^93^ with ggplot2^94^. For 3D analysis, images were analyzed in Imaris (Bitplane) for Tom20-DRP1 colocalization. Background subtraction was applied using a 10 µm filter, then surface renderings were generated of the Tom20 and DRP1 channels. Using the object-to-object statistics parameter, only DRP1 that was touching the Tom20 surface (0.0 µm away) was filtered for, and a separate surface channel was created to only include those DRP1 surfaces. Percent colocalization per cell was determined by taking the number of colocalized DRP1 surfaces divided by the total number of DRP1 surfaces in the cell, multiplied by 100.

### Mitochondrial membrane potential imaging and analysis

Cells were plated into a black 24-well plate (ibidi; Cat# 82426) and allowed to adhere overnight. Cells were stained with 100 nM MitoTracker Red CMXRos (ThermoScientific; Cat#: M7512) for 30 minutes at 37℃ and 5% CO2 and washed twice with 1X PBS. RPMI phenol red-free media containing 10% FBS and 1% P/S was then added into each well of the 24-well plate and cells were imaged on an EVOS M7000 high content imager (ThermoScientific) with 37℃ and 5% CO2 incubation. Images were analyzed in CellProfiler^95^ for mean fluorescence intensity of MitoTracker Red CMXRos staining.

### Transmission electron microscopy and mitochondrial cristae morphology analysis

Transmission electron microscopy sample prep was performed with a previously published protocol from Kostecka et al.^96^. Briefly, cells were cultured to the desired timepoints and were fixed overnight at 4℃ in a solution of 2.5% glutaraldehyde, 3mM MgCl2, in 0.1 M sodium cacodylate buffer, pH 7.2. Samples were rinsed with buffer followed by additional fixation in the dark and on ice for 1 hour with 1% osmium tetroxide reduced with 1.5% potassium ferrocyanide in 0.1 M sodium cacodylate buffer. Samples were rinsed with dH2O rinse and en bloc staining was performed for one hour in 2% uranyl acetate (aq). A graded series of ethanol was used to dehydrate the samples and samples were then embedded in Eponate resin overnight at 60℃. A diamond knife on the Reichert-Jung Ultracut E ultramicrotome was used to cut 60 to 90 nm sections. 2×1 mm formvar coated copper slot grids were used to pick up the sections. 2% uranyl acetate (aq.) and 0.4% lead citrate were used to stain the grids before imaging on a ThermoFisher Talos L120C at 120 kV with a ThermoFisher Ceta CCD (16 megapixel CMOS, 16-bit).

### Time lapse microscopy for mitochondrial dynamics

Cells were plated into 8-well ibiTreat µ-slides (ibidi; Cat#: 80826) and allowed to adhere overnight. Cells were stained with 150 nM MitoTracker Green (ThermoScientific; Cat#: M7514) for 30 minutes at 37℃ and 5% CO2 and washed twice with 1X PBS. RPMI phenol red-free media containing 10% FBS and 1% P/S was then added into each well of the 8-well ibiTreat µ-slide. Cells were imaged on a Zeiss LSM 880 microscope (Carl Zeiss) in Airyscan SR mode with an 63X, numerical aperture 1.4 PlanApo oil objective, at 37℃ and humidified 5% CO2. Cells were excited with a 488 nm laser and 2D images were taken every 30 seconds for 15 minutes at a resolution of 1280X1280 pixels to observe mitochondrial fission and fusion events. 2D Airyscan Processing was performed in Zen software, and images were then subjected to background subtraction in ImageJ^91^ using the rolling ball method with a radius of 50 pixels. Mitochondrial fission and fusion events were identified through Mitometer, a MATLAB-based software to track mitochondria over time^97^.

### Western blot

PC3 control cells and cells 10 Days PTR were lifted from T150 flasks with TrypLE^TM^ Express (GibcoTM; Ref# 12604-013) in lysate form using a RIPA buffer (Sigma-Aldrich; Cat#: R0278-500ML) with Halt Protease & Phosphatase cocktail (ThermoScientific; Cat#: 78440) and 0.5M EDTA (ThermoScientific; Cat#: 1861274). Lysates were incubated at 4℃ on a rotator for 30 minutes and then were subsequently spun in a 4℃ centrifuge at 21,000 rcf and stored at −20℃. The Pierce BCA Protein Assay was used to equally allocate 50 µg protein from each lysate for each lane being run in the gel. Each sample was made with the appropriate amount of lysate, 5 µL 4X Loading Buffer (4X Laemmli Buffer (Bio-Rad #1610747), beta-Mercaptoethanol (Bio-Rad; Cat#: 1610710XTU), and completed with UltraPure distilled water (ThermoScientific; Cat#: 10977023). Samples were boiled in a thermocycler set to 99℃ with a lid temperature of 105℃ for 10 minutes. A 4-20% Mini-PROTEAN TGX gel (Bio-Rad #4561094) was loaded with each sample and Chameleon Duo Pre-stained Protein Ladder (LI-COR Biosciences; Cat#: 928-60000). The gel run was set at 140 V for 60 minutes. For OPA1 probing, a 7.5% Mini-PROTEAN TGX gel (Bio-Rad #4561023) was used, and gels were run at 50 V for 300 minutes. Gels were transferred onto a 0.2 µm nitrocellulose membrane in the Trans-Blot Turbo Transfer Pack (Bio-Rad; Cat#: 1704158) using the BioRad Trans-Blot Turbo Transfer System. The membrane was blocked in 1X Casein Buffer (10X Casein Buffer (Sigma-Aldrich; Cat#: B6429-500ML) diluted with UltraPure distilled water on a room temperature shaker for 1 hour. The membrane was then incubated in a primary antibody specific to the target protein at a 1:1000 dilution in 1X Casein Buffer with mouse β-actin primary antibody (Sigma Aldrich; Cat#: A5441-0.5ML) at 1:5000 dilution overnight at 4℃. The primary antibodies used were Phospho-DRP1 Ser616 (Cell Signaling Technology; Cat# 3455S), Phospho-DRP1 Ser637 (Cell Signaling Technology; Cat#: 4867S), DRP1 (Cell Signaling Technology; Cat#: 8570S), MFF (Cell Signaling Technology; Cat#: 84580S), MiD49 (Proteintech; Cat# 28718-1-AP), MiD51 (Proteintech; Cat#: 20164-1-AP), OPA1 (Cell Signaling Technology; Cat#: 80471S), OMA1 (Cell Signaling Technology; Cat#: 95473S), MFN1 (Cell Signaling Technology; Cat#: 14739S), MFN2 (Cell Signaling Technology; Cat#: 9482S). For the OPA1 blots, Vinculin was used as the loading control (Cell Signaling; Cat#: 13901S) at a 1:1000 dilution. The following day, the membrane was washed four times for 5 minutes each in 0.1% TBS-Tween 20 (UltraPure distilled water, 10X TBS (Quality Biological #351-086-101CS), 0.1% Tween 20 (Sigma-Aldrich #9005645)). Then, the membrane was incubated in a secondary antibody diluted in 1X Casein Buffer for 1 hour on a room temperature shaker. The secondary antibodies used were IRdye® 800CW goat anti-rabbit IgG at a 1:15000 dilution, and IRdye® 680RD goat anti-mouse IgG at a 1:20000 dilution (LI-COR Biosciences; Cat# 926-32211 and 926-68070, respectively). The membrane was washed another four times for 5 minutes each in 0.1% TBS-Tween 20. Imaging was performed on the Azure Sapphire. (Azure Biosystems).

### Oxygen consumption measurements

Cells were cultured to the desired timepoints and lifted with TrypLE^TM^ Express (Gibco^TM^; Ref# 12604-013). TrypLE was neutralized with FBS-containing media and spun down at 500g for 10 minutes at 4 ℃. Cells were resuspended in 1 mL of complete RPMI media and put on ice until they were ready for oxygen consumption measurements. Oxygen consumption rate (OCR) of untreated cells and cells 10 Days PTR was measured over 10 minutes using the OxyTherm System Clark Type Oxygen Electrode (Hansatech Instruments). After measurements, cells were spun down, and cell weights were determined. OCR data were normalized to cell weight for accurate comparison between untreated cells and cells 10 Days PTR and plotted as nmol per mg of cell weight per minute.

### ATP/ADP Assay

Cells were cultured to the desired timepoints and lifted with TrypLE^TM^ Express (Gibco^TM^; Ref# 12604-013). TrypLE was neutralized with FBS-containing media and spun down at 500g for 10 minutes at 4 ℃. Cells were resuspended in 1 mL of complete RPMI media and put on ice until they were ready for lysis. Cells were adjusted to a concentration of 10,000 cells per 10 µL for untreated cells and a concentration of 4,000 cells per 10 µL for cells 10 Days PTR. Cells were lysed with 90 µL of Assay Buffer from the ATP/ADP Assay Kit (Sigma Aldrich; Cat#: MAK135). ATP detecting reagents were mixed with the cell lysate and luminescence was measured with the FLUOstar Omega plate reader (BMG LABTECH). ADP converting enzyme was then added into the mixture to convert ADP to ATP for a second luminescence read as a surrogate for ADP levels.

### siRNA transfection

The Mission siRNA Universal Negative Control (Sigma Aldrich; Cat# SIC001) and *Oma1* siRNA sequences (Sigma Aldrich; Cat#: SASI_Hs01_00136476 & SASI_Hs01_00136477) are indicated as Scramble, *Oma1* siRNA#1 and *Oma1* siRNA#2, respectively. Their sequences can be found in **Table 1**. Cells were plated with RPMI media containing 10% FBS without antibiotics into either black Nunc^TM^ 96-well plates (ThermoScientific; Cat#: 165305) or T150 flasks (Corning; Cat# 430825) and allowed to adhere overnight. The Mission siRNA Universal Negative Control, *Oma1* siRNA#1, and *Oma1* siRNA#2 siRNA were diluted in Opti-MEM^TM^ I Reduced Serum Media (ThermoScientific; Cat#: 31985070). Lipofectamine^TM^ RNAimax transfection reagent (ThermoScientific; Cat#: 13778075) was diluted in Opti-MEM^TM^ I Reduced Serum Media. The diluted siRNAs and Lipofectamine^TM^ RNAimax were mixed together at a 1:1 ratio and incubated at room temperature for 20 minutes. The siRNA-Lipofectamine complexes were added onto cells cultured in Opti-MEM^TM^ I Reduced Serum Media to obtain a final siRNA concentration of 50 nM. Cells were incubated at 37℃ and 5% CO_2_ for 72 hours and assayed for cell viability or harvested for protein lysates.

**Table 1.**
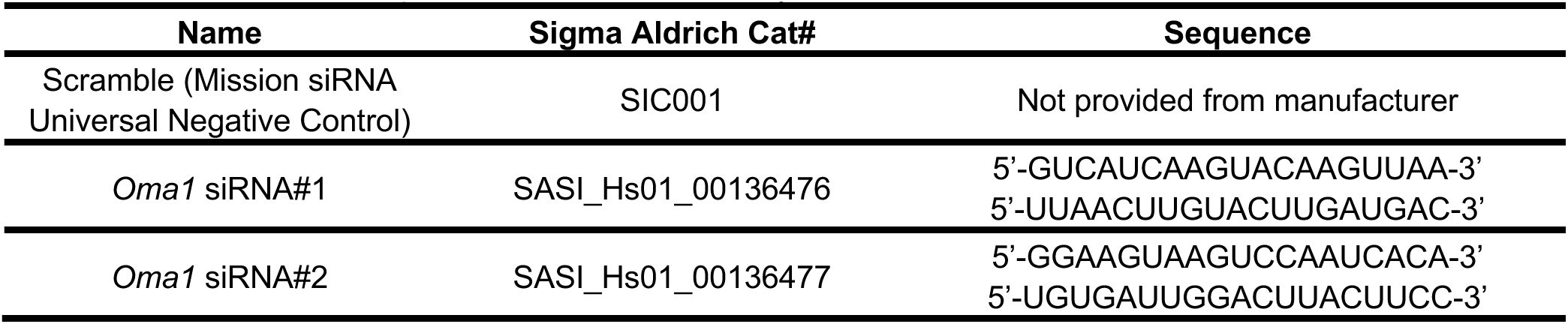
*Oma1* siRNA sequences used in this study.

### Cell viability assay

Cells were plated in black Nunc^TM^ 96-well plates (ThermoScientific; Cat#: 165305) and allowed to adhere overnight. Cells were transfected with *Oma1* siRNA and Scramble negative control for 72 hours. At endpoint, the transfection medium was removed, and cells were washed with 1X PBS. Cells were then stained for 20 minutes with 5 µg/mL Hoechst 33342 (ThermoScientific; Cat#: H3570) diluted in RPMI phenol red-free media containing 10% FBS and 1% P/S. Following staining, cells were imaged immediately for DAPI signal with the EVOS M7000 high content imager (ThermoScientific) with 37℃ and 5% CO2 incubation.

Images were imported into ImageJ (NIH) and nuclei counts were obtained with a custom batch processing ImageJ script. Briefly, background subtraction was performed using the rolling ball method with a radius of 50 pixels. The 16-bit image was converted to 8-bit, and a threshold was set for Hoechst signal. To count the thresholded nuclei, the “Analyze Particles” command was used. Nuclear counts were used as a proxy for viable cell count. Data was plotted as percent cell viability normalized to Scramble negative control.

### Single cell RNA sequencing analysis

The transcriptome of untreated cells and Cells 10 Days PTR was deposited previously by our group into the Gene Expression Omnibus (GEO) database (GSE297299)^98^. Data was downloaded and imported into the Partek Flow software (Illumina) for normalization, pseudo-bulking, and downstream analysis. Gene set enrichment analysis (GSEA) was performed in Partek Flow with HALLMARK gene sets extracted from the Molecular Signature Database (MSigDB)^99,100^.

### Statistics

Statistical analyses were performed in GraphPad Prism version 9.1 (GraphPad Software, LLC) and RStudio^93^. Statistical tests performed are reported in respective figure legends. An alpha value of 0.05 was used for all tests. ns, *, **, ***, and **** indicated p>0.05, p<0.05, p<0.01, p<0.001, and p<0.0001, respectively. In the violin plots, the solid line represents the median, and the dotted lines represent the lower and upper quartiles.

## DATA AVAILABILITY

Data is provided within the manuscript or in supplementary information files.

## Supporting information

Supplementary Information_Updated

## ACKNOWLEDGEMENTS

This work was supported by the William and Carolyn Stutt Research Fund, Ronald Rose, MC Dean, Inc., William and Marjorie Springer, Mary and Dave Stevens, Louis Dorfman, and the Jones Family Foundation. We thank the Johns Hopkins School of Medicine Microscope Facility for the acquisition of transmission electron microscopy images with the ThermoFisher Talos L120C. We also thank Dr. Hiromi Sesaki and Dr. Anne Hamacher-Brady for helpful discussions throughout this project.

## FUNDING

This work was supported by the US Department of Defense CDMRP/PCRP 367 (W81XWH-20-10353), the Prostate Cancer Foundation, and the Patrick C. Walsh Prostate Cancer Research Fund to SRA; and NCI grants U54CA143803, CA163124, CA093900, and CA143055, and the Prostate Cancer Foundation to KJP. This work was also supported by the Office of the Director and the National Institute of General Medical Sciences of the National Institutes of Health under award number S10OD023548.

## AUTHOR CONTRIBUTIONS

Conceptualization: ML, KJP, SRA; Experimental work: ML, CAM, LTAR, QN, ZG; Data analysis and figure generation: ML; Writing – original draft preparation: ML; Writing – review and editing: ML, KJP, SRA. Funding acquisition: SXS, KJP, SRA.

## COMPETING INTERESTS

KJP is a consultant to Cue Biopharma, Inc., an equity holder in PEEL Therapeutics, and a founder and equity holder in Keystone Biopharma, Inc. SRA is an equity holder in Keystone Biopharma, Inc. The companies were not involved in the design, collection, analyses or interpretation of data, writing of the manuscript, or the decision to publish the results.

## ETHICS DECLARATIONS

Approval for animal experiments: Not applicable

Approval for human experiments: Not applicable

Consent to participate/consent to publish: Not applicable

## SUPPLEMENTARY INFORMATION

**Supplementary Fig. S1.** N-acetyl cysteine decreases levels of reactive oxygen species in cells 10 Days Post-Treatment Removal. (a) Representative DCF-DA fluorescence images of cells treated with N-acetyl cysteine and vehicle. (b) Quantification of mean fluorescence intensity of DCF-DA between indicated groups.

**Supplementary Fig. S2.** Glycolysis and hypoxia gene sets are enriched in cells 10 Days PTR. Gene set enrichment analyses from single-cell RNA sequencing data of cells 10 Days PTR and untreated cells. (a) Enrichment plot of the HALLMARK GLYCOLYSIS gene set (b) Enrichment plot of the HALLMARK HYPOXIA gene set.

**Supplementary Fig. S3.** Uncropped blot images from Figure 2a. Boxed areas delineate the cropped portion displayed in Figure 2a.

**Supplementary Fig. S4.** Uncropped blot images from Figure 3a. Boxed areas delineate the cropped portion displayed in Figure 3a.

**Supplementary Fig. S5.** Uncropped blot images from Figure 5a. Boxed areas delineate the cropped portion displayed in Figure 5a.

**Supplementary Video 1.** Representative time lapse of mitochondrial dynamics in untreated PC3 cells stained with MitoTracker Green.

**Supplementary Video 2.** Second representative time lapse of mitochondrial dynamics in untreated PC3 cells stained with MitoTracker Green.

**Supplementary Video 3.** Representative time lapse of mitochondrial dynamics in cells 10 Days Post-Treatment Removal stained with MitoTracker Green.

**Supplementary Video 4.** Second representative time lapse of mitochondrial dynamics in cells 10 Days Post-Treatment Removal stained with MitoTracker Green.

## Notes

### Summary of Updates

Figure 3 revised; Figure 5 revised; Figure 6 added. Supplemental files updated; Results and Discussion sections updated to report on revised figures.

