## Supplementary Information_Updated for "Cancer cells surviving cisplatin chemotherapy increase stress-induced OMA1 activity and mitochondrial fragmentation"

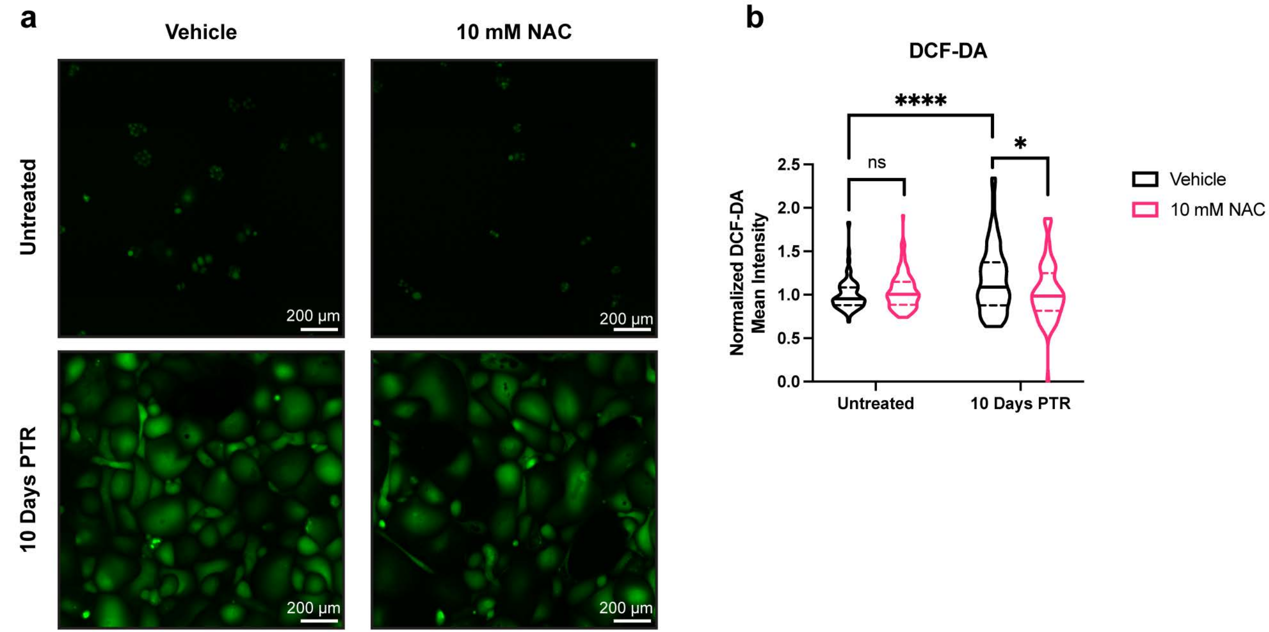


**Supplementary Fig. S1. N-acetyl cysteine decreases levels of reactive oxygen species in cells 10 Days Post-Treatment Removal.** (**a**) Representative DCF-DA fluorescence images of cells treated with N-acetyl cysteine and vehicle. (**b**) Quantification of mean fluorescence intensity of DCF-DA between indicated groups.

**Supplementary Fig. S2. Glycolysis and hypoxia gene sets are enriched in cells 10 Days PTR.** Gene set enrichment analyses from single-cell RNA sequencing data of cells 10 Days PTR and untreated cells. (**a**) Enrichment plot of the HALLMARK GLYCOLYSIS gene set. (**b**) Enrichment plot of the HALLMARK HYPOXIA gene set.

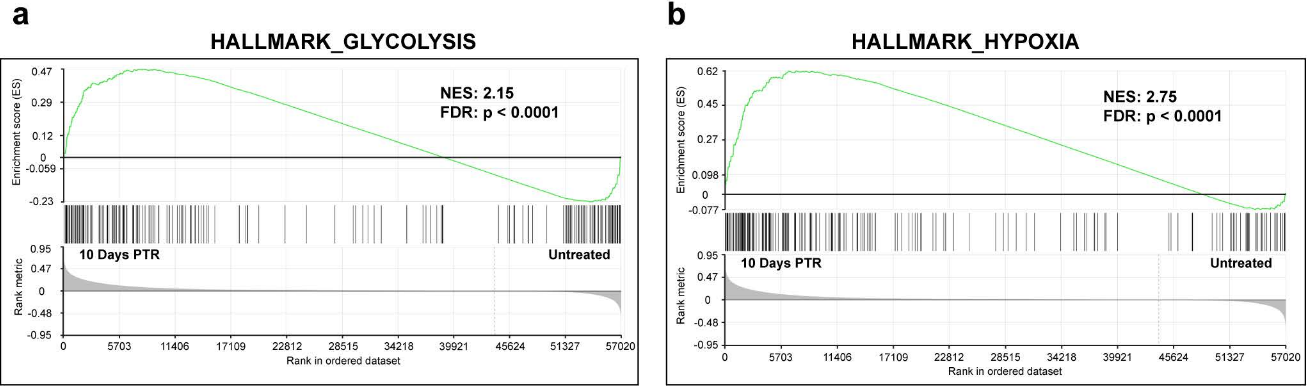

**Supplementary Fig. S3. Uncropped blot images from Figure 2a.** Boxed areas delineate the cropped portion displayed in Figure 2a.

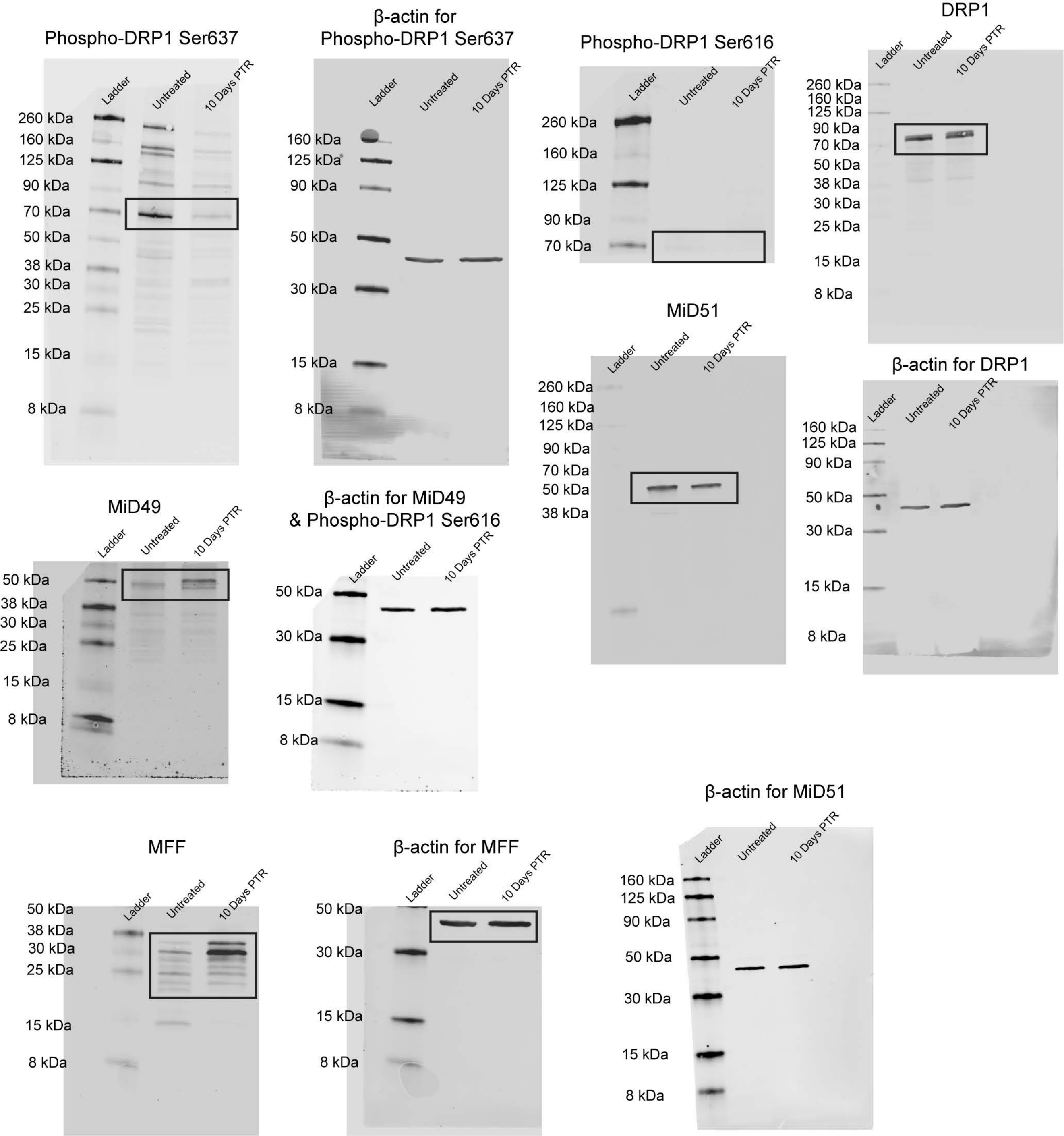

**Supplementary Fig. S4. Uncropped blot images from Figure 3a.** Boxed areas delineate the cropped portion displayed in Figure 3a.

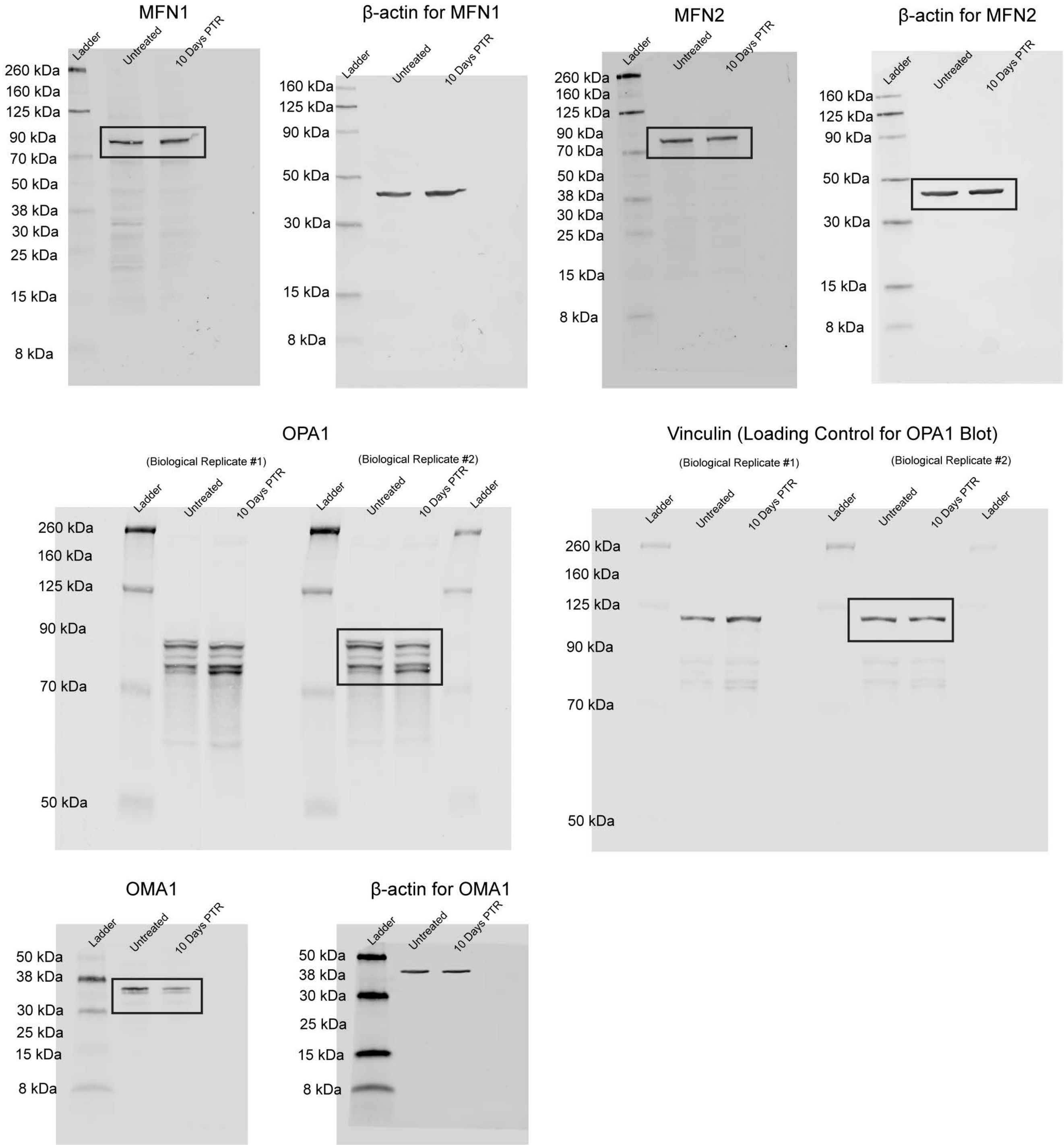

**Supplementary Fig. S5. Uncropped blot images from Figure 5a.** Boxed areas delineate the cropped portion displayed in Figure 5a.

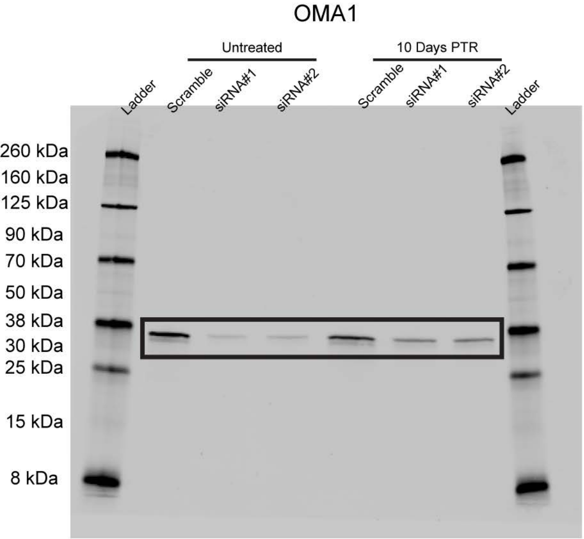


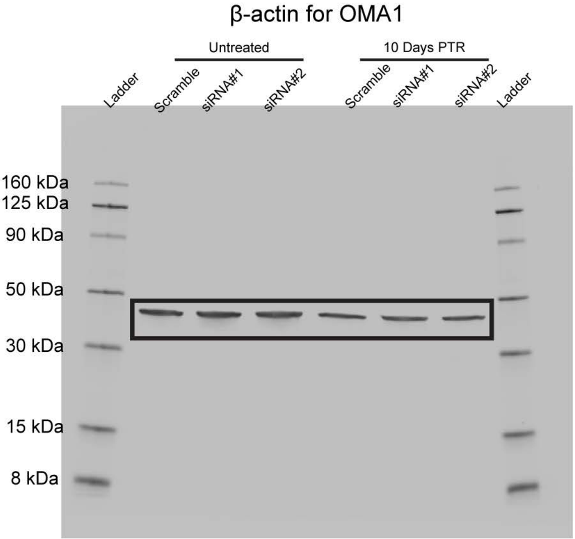

**Supplementary Videos 1-4.**

Due to the video size exceeding bioRxiv's 40 MB limit, the supplementary videos are deposited in a OneDrive Folder and can be viewed through the link below.

https://livejohnshopkins-my.sharepoint.com/:f:/g/personal/mli154_jh_edu/EkQbnbLWvZhKu1v6M18h5VoBo30PW1Ie-EjUgUkBh7vBtw
